# PAV-spotter: using signal cross-correlations to identify Presence/Absence Variation in target capture data

**DOI:** 10.1101/2024.10.25.620064

**Authors:** Manon de Visser, Chris van der Ploeg, Milena Cvijanović, Tijana Vučić, Anagnostis Theodoropoulos, Ben Wielstra

## Abstract

High throughput sequencing technologies have become essential in the fields of evolutionary biology and genomics. When dealing with non-model organisms or genomic gigantism, sequencing whole genomes is still relatively costly and therefore reduced-genome representations are frequently obtained, for instance by ‘target capture’ approaches. While computational tools exist that can handle target capture data and identify small-scale variants such as single nucleotide polymorphisms and micro-indels, options to identify large scale structural variants are limited. To meet this need, we introduce PAV-spotter: a tool that can identify presence/absence variation (PAV) in target capture data. PAV-spotter conducts a signal cross-correlation calculation, in which the distribution of read counts per target between samples of different *a priori* defined classes – e.g. male versus female, or diseased versus healthy – are compared. We test, validate and apply our methodology by studying *Triturus* newts: salamanders with gigantic genomes that currently lack an annotated reference genome. *Triturus* newts suffer from a hereditary disease that kills half their offspring during embryogenesis. We compare the target capture data of two different types of diseased embryos, characterized by unique deletions, with those of healthy embryos. Our reciprocal, self-validating findings show that PAV-spotter helps to expose such structural variants, even in the face of medium to low sequencing coverage levels, low sample sizes, and background noise due to mis-mapped reads. PAV-spotter can be used to study the structural variation underlying supergene systems in the absence of whole genome assemblies. The code, including further explanation, is available through the PAV-spotter GitHub repository: https://github.com/Wielstra-Lab/PAVspotter.

## Introduction

Next-generation DNA sequencing methods have revolutionized the biological sciences, with an ever-growing amount of sequence data being generated worldwide (Annavi et al, 2019; Morganti et al, 2019). High throughput sequencing techniques have become more affordable and increasingly used, however sequencing entire genomes can still be challenging, for instance when dealing with non-model organisms, genomic gigantism, or a combination of the two (Ekblom & Galindo, 2011; Lemmon & Lemmon, 2013; Wachi et al, 2018). In such cases, well-annotated reference genomes for aligning (re-)sequenced reads are generally unavailable, making whole genome sequencing relatively costly in terms of money and (computational) time (e.g. see Etherington et al, 2020; Rovelli et al, 2019; Zaharias et al, 2020). Many biologists therefore still opt for more cheap and efficient ‘reduced-representation’ high throughput sequencing techniques, which allow for a subset of loci to be sequenced more deeply (Andermann et al, 2019; Da Fonseca et al, 2016; Puritz & Lotterhos, 2018).

A technique that has become particularly popular for studying non-model species is target capture, also referred to as hybridization sequencing, exon capture, sequence capture, or exome capture (Bi et al, 2012; Kaur & Gaikwad, 2017; Puritz & Lotterhos, 2018; Yohe et al, 2020). This method facilitates collecting sequence information on hundreds or thousands of pre-selected loci. Due to the consequent rise of large multi-locus DNA sequence datasets, the need for innovative, easy-to-implement bioinformatic applications has surged as well (Gauthier et al, 2019). User-friendly pipelines help to pre-process sequence reads by wrapping and connecting existing software tools, with well-known examples for target capture including HybPiper (Johnson et al, 2016), Assexon (Yuan et al, 2019), Sequence Capture Processor (i.e. SECAPR, see Andermann et al, 2018), and CAPTUS (Ortiz et al., 2023). These pipelines and their software dependencies support fast upstream data cleaning and guide the user to phylogenetic applications downstream.

Typically, target capture analysis pipelines are utilized to identify small-scale genetic variants such as single nucleotide polymorphisms (SNPs) and relatively small insertions/deletions (microindels, ranging from 1∼50bp) to perform, for instance, phylogenetic tree-building (Andermann et al, 2019). However, to identify and analyze larger scale information from target capture data, such as genomic structural variation, few tools are available – especially when focusing on non-model organisms. A particular type of larger-scale variation that is hard to identify in multi-locus DNA sequence datasets is presence/absence variation (hereafter referred to as ‘PAV’) of relatively big insertions/deletions (macroindels, > 50bp), i.e. above the size of microindels (Wang et al, 2014; Zhang et al, 2014).

Structural variants such as PAVs are regularly overlooked because they are harder to identify than SNPs (Mahmoud et al, 2019; Wold et al, 2021). However they are a major source of genetic divergence and diversity (Gerdol et al, 2020; Hu et al, 2022; Marroni et al, 2014; Rosa et al, 2015). PAV in particular poses an extreme example of copy number variation, where fragments in the size of entire exons or (stretches of DNA containing multiple) genes are missing from one genome compared to another (Gabur et al, 2020; Wellenreuther et al, 2019). When comparing such genomes, target capture data would in theory display PAV by showing ‘normal data’ in the case of target presence, versus a ‘data gap’ in the case of target absence. This would be an indication of structural variation.

Whether there are consistent differences in presence or absence of sequence data can be determined by analyzing the way that reads pile up against a certain reference set of sequences. Some tools can detect copy number variation and PAV patterns by comparing the depth of mapped reads of different samples, such as ExomeCNV (Sathirapongsasuti et al, 2011) and SUPER-CAP (Yuan et al, 2019). However, these tools come with strict requirements, such as good quality reference genomes being available, known functional annotation of variants, and/or coverage being consistently high across all samples and targets, with mapped reads ideally following a normal distribution. Yet, most multi-locus datasets, including those resulting from target capture experiments, generally do not meet such requirements (Andermann et al, 2019).

We introduce PAV-spotter: a flexible signal cross-correlation method that is able to ‘spot’ potential PAV in target capture datasets. Our approach borrows the notion of cross-correlation to detect the dissimilarity between datasets obtained through target capture experiments. Cross-correlation methods are generally used in the domain of control engineering. Classically, they are applied on time-series data (Verhaegen & Verdult, 2007), for example in machinery fault detection studies (Gao et al, 2015). However, cross-correlation approaches have been proven useful in the field of pattern recognition as well (Jain et al, 2000).

Being able to identify structural variation by using pattern recognition would be especially informative when studying supergenes systems, in which individuals can have zero, one or two copies of particular loci (Hall & Wayne, 2013; Thompson & Jiggins, 2014). Supergenes consist of genes that are inherited together as a single locus due to the suppression of recombination (Berdan et al, 2022; Gutierrez-Valencia et al, 2021; Thompson & Jiggins, 2014). As a result, the non-recombining stretches of ‘supergene DNA’ evolve independently of one another, facilitating the rapid evolution of complex adaptations (Pennisi, 2017; Schwander et al, 2014). These sets of genes are often polymorphic and subjected to balancing selection, as a species generally possesses at least two supergene variants (Llaurens et al, 2017).

Sex chromosomes, for instance, are classically considered supergenes (Joron et al, 2006). In diploid organisms the heterogametic sex inherits the sex-determining ‘supergene’, as well as the alternate sex chromosome, in a hemizygous manner – meaning that they only receive one copy of each (Dufresnes & Crochet, 2022). In the XX-XY sex determination system of mammals, for instance, males generally possess a single copy of the supergene that is the Y chromosome (as well as a single copy of the X-chromosome), whereas in the ZW-WW system it is the females that possess the Z supergene once (next to a single W chromosome). Hence, genes that are hemizygous and thus lie solely on the sex-determining supergene would show PAV in target capture datasets when data of different sexes is compared.

However, supergene systems are not limited to the biological concept of sex. Other, famous examples of supergenes underlying complex traits are; the Müllerian mimicry complex in *Numata* longwing butterflies (Joron et al, 2006), the striking sexual dimorphism and breeding behaviors of ruffs (Kupper et al, 2016; Lamichhaney et al, 2016) and white-throated sparrows (Tuttle et al, 2016), the social polymorphism observed in several species of ant (Kay et al, 2022), and heterostyly in primrose flowers (Li et al, 2016). Furthermore, hemizygous inheritance of (super)genes also occurs in, for instance, genetic incompatibilities such as with “hybrid necrosis” in plants, which can be linked to PAV in certain genes in for example Asian rice (Li et al, 2023; Li & Lee, 2023); hereditary diseases such as α-thalassaemia, which is caused by large deletions in the alpha globin gene cluster on chromosome 16 in humans (Harteveld & Higgs, 2010); and in balanced lethal systems, in which two distinct chromosome forms exist that are covered by unique lethal mutations (Wielstra, 2020).

We demonstrate the application of PAV-spotter using the balanced lethal system in *Triturus* newts as a case study. *Triturus* individuals either are heteromorphic and possess two different versions of their largest autosomal chromosome, characterized by unique deletions, or they are homomorphic and possess two identical versions of this chromosome (De Visser et al, 2024). The two types of homomorphic individuals express a unique disease state and both die during embryogenesis, whereas the heteromorphic individuals are viable – a phenomenon caused by evidential PAV (De Visser et al, 2024; France et al, 2024; Macgregor & Horner, 1980; Meilink et al, 2025; Sessions et al, 1988; Wallace, 1987). This ‘double hemizygous’ system lends itself particularly well for using target capture data to detect PAV, as it allows for a reciprocal, self-validating test: targets deleted from one chromosome version should be present on the alternate version and the other way around. This approach, in which we apply the same analytical framework to both chromosome versions, is particularly valuable given the importance of methodological consistency in accurately detecting PAV (Brůna et al., 2026). Based on our findings, we describe the usefulness, as well as the limitations, of our tool.

## Methods

### Sample information & collection

The first chromosome of *Triturus* comes in two forms: 1A and 1B. Homomorphic individuals (1A1A or 1B1B) invariably die during embryogenesis, while heteromorphic individuals (1A1B/1B1A) are viable. We collected *T. macedonicus* x *T. ivanbureschi* F_1_ hybrid embryos from an ongoing breeding experiment at the Institute for biological research, „Siniša Stanković”, University of Belgrade (with experimental settings, breeding conditions, and other details on the process of raising embryos as described in Vucic et al, 2024; Vučić et al, 2022). Embryo development was followed through observation with a stereomicroscope. Diseased embryos were collected when the process leading up to developmental arrest occurred, which is visible as a ‘growth slowdown’, during the late tail-bud phase. Diseased embryos were then classified into either the “fat-tailed” (FT) phenotype or the “slim-tailed” (ST) phenotype based on morphological characteristics of the embryo (Sessions et al, 1988; Sims et al, 1984). Healthy/control (HC) embryos that survived this critical phase were subsequently collected. We collected 30 individuals in total (Supplementary Table 1); ten of each class, i.e. ten ST, ten FT and ten HC embryos. Samples were stored in ethanol at −20 °C until further handling.

### Laboratory procedures & pre-processing of sequence data

We followed the standard “NewtCap” workflow of salt-based extraction of DNA from embryonic tissue, followed by quantification, library preparation, target capture and Illumina sequencing (as described in De Visser et al, 2025a). After obtaining the raw, paired-end sequence reads from Baseclear B.V. (Leiden, the Netherlands), we followed a standard pipeline for checking the quality of, and for cleaning-up and mapping, our sequence data in a Linux environment up to and until the deduplication of the BAM files step (as described in De Visser et al, 2025a). These deduplicated BAM files served as input for further data extraction and analyses. Throughout the cleaning and mapping process, we used SAMtools’ (Danecek et al, 2021) *stats*, *flagstat* and *coverage* options to calculate basic statistics from the FASTQ and BAM files. The reference FASTA file used for read mapping can be found in Supplementary Material as ‘Targets.fasta’. These 7,139 sequences, initially used for probe tiling, were based on *T. dobrogicus* transcripts, and had a maximum length of 450bp (De Visser et al, 2025a; Wielstra et al, 2019).

### Preparing read depth data for PAV-spotter

From the BAM files we extracted information on sequence read depth for all sites per target by using the SAMtools depth option (Danecek et al, 2021). We optimized this extracted information by following several file-manipulation steps in a custom ‘prepping’ shell script – “Script1_prepping.sh” – to make the input files and folder structure match the requirements of our PAV-spotter tool. The steps in Script1_prepping.sh include automatically merging and sorting of intermediate files where appropriate, changing the tab-delimited format to a CSV format, splitting the overall CSV file into multiple files (one file per separate target/gene), and creating a text file with sample names for later use (details are explained in the script). The exact format of the input folder structure and input files is described in Box 1.

#### BOX 1

*This is a description of the expected input file format for the main PAV-spotter scripts:*

***<u>‘Species’ Directories:</u>***

*- Each species, or otherwise distinguishable set of data, has its own directory*
*- PAV-spotter is built in such a way that it will loop over multiple such directories*

***<u>“individuals.txt” file:</u>***

*- An automatically generated file, uses the initial sample names for input*
*- Needs to be located in, and corresponding to the contents of, a particular species directory*
*- This file contains the individual information, with each sample name on a new line and with information on the classes to be compared included in the name (e.g. ‘ST’, ‘FT’, and ‘HC’)*

***<u>Gene/Target Data Files:</u>***

*- Also needs to be located in, and corresponding to the contents of, a particular species directory*
*- Each file represents a single gene or target*
*- Each file has columns and is in CSV format (this should be the output from batch script 1 “Script1_prepping.sh”):*

*- Column 1: Gene/target name*
*- Column 2: Gene/target position (a number)*
*- Column 3: Read depth data (a number)*
*- Column 4: Sample/class name (should match identifiers in “individuals.txt” file names)*

PAV-spotter assumes background knowledge on phenotype classes that presumably differ in the presence of certain genes (in other words, cross-comparisons are not random: for instance males are compared to females, diseased individuals are compared to control samples, etc.). Here, we work with the a priori classification of three phenotypes: two types of diseased embryos (FT vs. ST) and healthy embryos to serve as a control (HC). Which sample belongs to which class needs to be designated by your input filenames (the BAM files), which will end up in an automatically created text file “individuals.txt” after running Script1_prepping.sh. This is crucial, as PAV-spotter performs the comparisons based on the phenotype information embedded in the names of this text file. In case phenotypic classes as specified by the user cannot be deduced from input file names, the user needs to either alter the input file names manually, or alter the identifiers in Script1_prepping.sh manually – or both – before running Script1_prepping.sh (ideally, the user includes such filename identifiers already at the raw FASTQ file stage for consistency throughout the entire bioinformatics procedure).

### Applying PAV-spotter

We applied Script1_prepping.sh on our total set of 30 samples (Supplementary Table 2; n=30, ten ‘ST’, ten ‘FT’ and ten ‘HC’ individuals, with these identifier abbreviations occurring in the sample names). Additionally, we applied the script on random subsets of samples (Supplementary Table 3 and Supplementary Table 4); two separate analyses with a sample size of five per class (i.e. two total subsets of n=15, indicated by sample set ‘5_1’ and run ‘5_2’), and five more separate analyses with a sample size of two per class (i.e. five total subsets of n=6, indicated by sample set ‘2_1’, ‘2_2’, ‘2_3’, ‘2_4’ and ‘2_5’). This allowed us to assess the performance of PAV-spotter when lower sample sizes are used. We randomized the grouping of samples into subsets by using the ‘shuf’ command from the standard GNU Core Utilities (http://gnu.org/s/coreutils/).

We ran a custom MATLAB script (“Script2b_PAVSPotter.m”, hereafter ‘PAV-spotter’ – also available in Python language as “Script2b_PAVSpotter.py”) remotely through SLURM workload manager (example batch scripts attached as “Script2a_scheduler_matlab.sh” and “Script2a_scheduler_python.sh”, respectively). Users can compare two, or three phenotypic classes (using the argument “categories”, see the SLURM scripts), but if only two are provided, only two are compared. Also, we indicated which class we considered the control group (by implementing “ctrl_category = ‘HC’”) and we provided a common identifier for all input files/targets (with the argument “common_identifier = ‘DN’”). Also, information on the working directory and desired output filename was provided in the MATLAB command depending on the analysis, and the same is the case for the customized #SBATCH lines for running the SLURM job.

PAV-spotter was run four times on our total set of n=30: once with default settings (no filtering, “reads_threshold” == 0 and “contig_width” == 0), one time with a mild coverage filtering (“reads_threshold” == 5 and “contig_width” == 0), one time with a mild filtering for minimum length of contigs (“reads_threshold” == 0, “contig_width” == 50) and one time with both of the filtering thresholds (“reads_threshold” == 5 and “contig_width” == 50). By setting a soft coverage filter of a minimum of five reads, we filter out reads of any poorly covered target of an individual that does not meet this criterium (thereby assuming a coverage of zero across the target in question). Furthermore, by specifying a minimum contig width, the script will filter out read information of covered regions in between two positions with zero coverage, in case those regions are narrower than, in this case, 50bp (thereby assuming a coverage of zero in that specific target region). For the tests on sample subsets of five individuals per class and two individuals per class, we only used the default filtering settings (no filtering, “reads_threshold” == 0 and “contig_width” == 0). In all analyses, we enabled the script to plot accessory figures (“plot_figures” == TRUE, clarification below). We ran PAV-spotter using MATLAB v.9.13.0 (https://www.mathworks.com) for the main components of our overall study and added the Python version of the tool in order to make it more widely accessible. We used Python v.3.11 to run tests and to ensure identical performance and output files.

### The rationale behind PAV-spotter

After setting up the input files and initiating PAV-spotter successfully, filtering settings are applied as specified. Subsequently, PAV-spotter merges and normalizes the sequence read depth information for all the samples per target per specified sampling class before it starts the actual cross-correlation analyses (but users can turn this setting off in case calculations of all possible cross-comparisons on an individual level are desired). By exploiting the availability of multiple samples per phenotypic class in this way, we ensure that the comparisons will be made on a class level rather than on an individual level. In the latter case, more false outcomes would be expected as a result of the stochastic nature of target capture experiments, something that can be avoided by pooling results. PAV-spotter can loop through a set of input directories if separate datasets need to be analyzed with similar settings consecutively.

The cross-correlation analysis in PAV-spotter works as follows (Figure 1). Two target capture results of the same targeted region, but of a different phenotypic class, are defined by D_1_and *D*_2_. These represent two vectors of identical length *T*, where the vector position represents the target position and the vector values represent the normalized sequence read depth. PAV-spotter then calculates the similarity of the cross-correlation formulated by

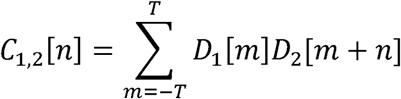

for all *n* between *-T* and *T*, with *n* and *m* accessing the indices of the vectors *C*_1,2_, *D*_1_, *D*_2_. This constructs a cross-correlation vector of which the maximum value provides the maximum similarity of *D*_1_and *D*_2_. For example, in the case that *D_1_* = *D_2_* (i.e., in the case of autocorrelation), the similarity score C will be 100%.

**Figure 1:**
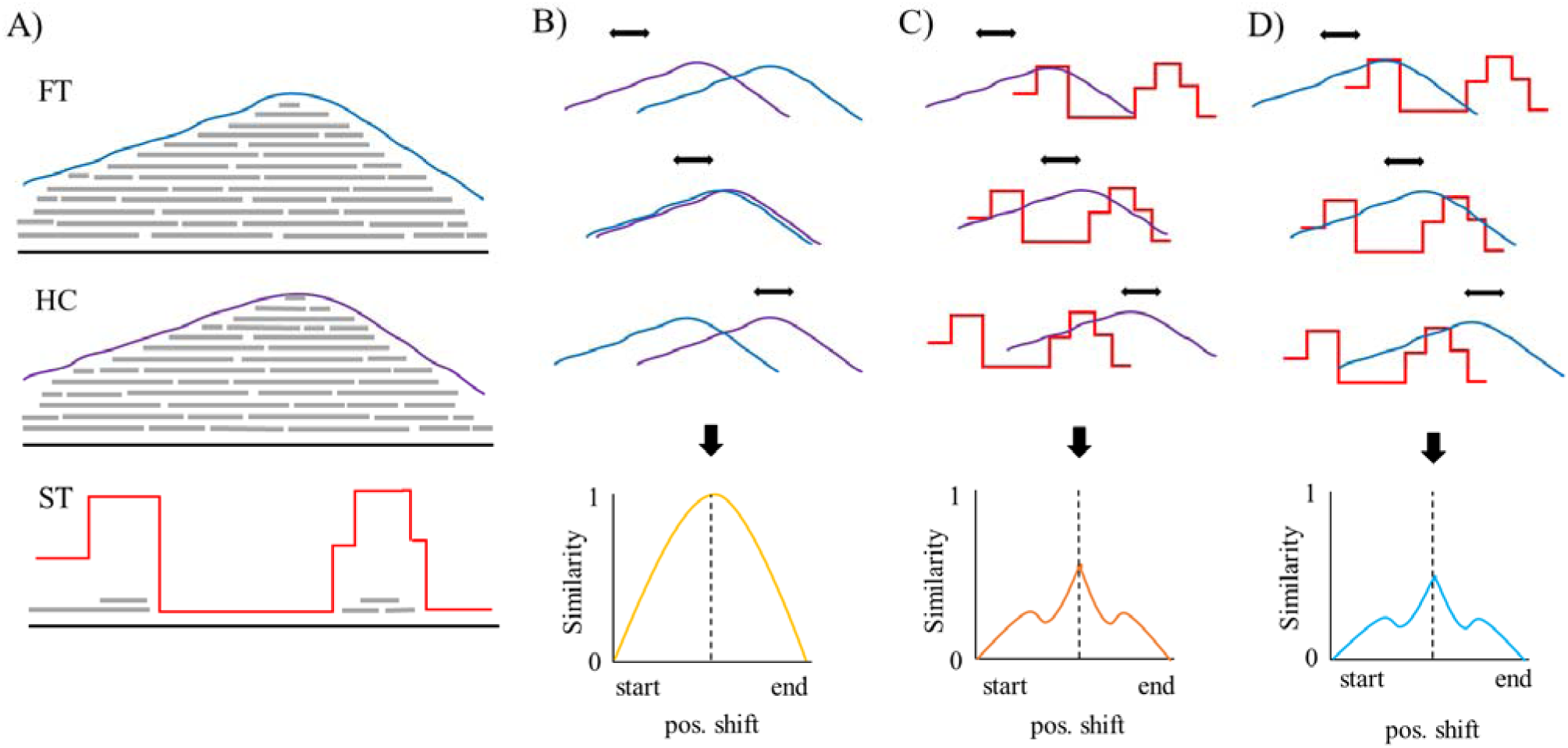
A simplified visualization of the cross-correlation methodology in PAV-spotter, showing a hypothetical gene/target that is missing in only ST embryos as an example. **(A)** A number of sequence reads are mapped (in grey) against a gene/target. This information is taken from depth files and merged per phenotypic class (in case of multiple samples per class), then the absolute distributions of the read depths are normalized, here represented by the colored lines; dark blue = FT, purple = HC, red = ST. **(B)** The distribution data of the FT and the HC classes are compared. A measure of similarity is determined as a function of the displacement of one read depth distribution relative to the other, as if they were to ‘slide over’ each other (indicated by black arrows). The similarity appears close to 1 (=100%) and the graph produced by PAV-spotter also follows a smooth line. **(C)** The distribution data of the HC and ST classes are compared: the similarity is not close to 100% and the similarity graph produced by PAV-spotter does not follow a smooth line. **(D)** The distribution data of the HC and ST classes are compared: the similarity is not close to 100% and the similarity graph produced by PAV-spotter does not follow a smooth line.

PAV-spotter outputs a CSV file per separate analysis with all the cross-correlation data in the form of percentage similarity, and it outputs a folder with figures. Also, a CSV file with cross-comparisons between data from samples from the control group only (the HCs) is generated. This extra file allows for a calculation of the overall resemblance of the control samples, which should have data present with well-captured targets. This as opposed to the FT and ST samples, which are expected to show ‘data gaps’, or absence, for some targets.

As we are investigating a double hemizygous system, we always have three cross-correlation values to work with. This means we are able to use not only the healthy embryos (HC), but also the other class of diseased embryo (ST or FT), as a control, because genes absent in one class of diseased embryo are expected to be present in both other embryo classes (e.g. to recognize absence in ST embryos, which should have a 1A1A genotype, we can check for presence in the HC embryos which should have the 1A1B genotype, but we can do an additional check for presence in the ST embryos that should have a 1B1B genotype – and the same applies the other way around). Hence, to deduce PAV in the chromosome that is inherited twice in ST embryos, we search for a pattern in which a significant portion of the target was present in both HC and FT embryos (which contain the alternate chromosome form), but absent in ST embryos. Conversely, to deduce PAV in the chromosome inherited twice by FT embryos, we searched for absence in FT, but presence in both HC and ST embryos.

### Downstream PAV estimation

To automatically deduce PAV patterns, we applied another custom shell script: “Script3_downstream.sh”. This script takes the main output file of PAV-spotter, creates an overall matrix of the results, adds columns with information on the cross-correlation data that stand out, and makes lists of the targets that show potential PAV based on a threshold (which can be customized). When, for a certain target, the data of FT versus ST embryos were less than 80% similar, the data of FT versus HC embryos were less than 80% similar, and the data of ST versus HC embryos were more than 80% similar, this target was scored as ‘1A-linked’. For ‘1B-linked’ genes it was the other way around: FT vs. ST embryos < 80% similar, ST vs. HC embryos < 80% similar, and FT vs ST embryos > 80% similar. This threshold of 20% dissimilarity equals a p-value of between 0.01 (≈ 15% dissimilarity) and 0.001 (≈ 28% dissimilarity). The PAV-spotter output file that shows all similarity scores among the HC embryos only, guided our choice for this threshold (e.g. see Supplementary Figures 1 and 2).

**Figure 2:**
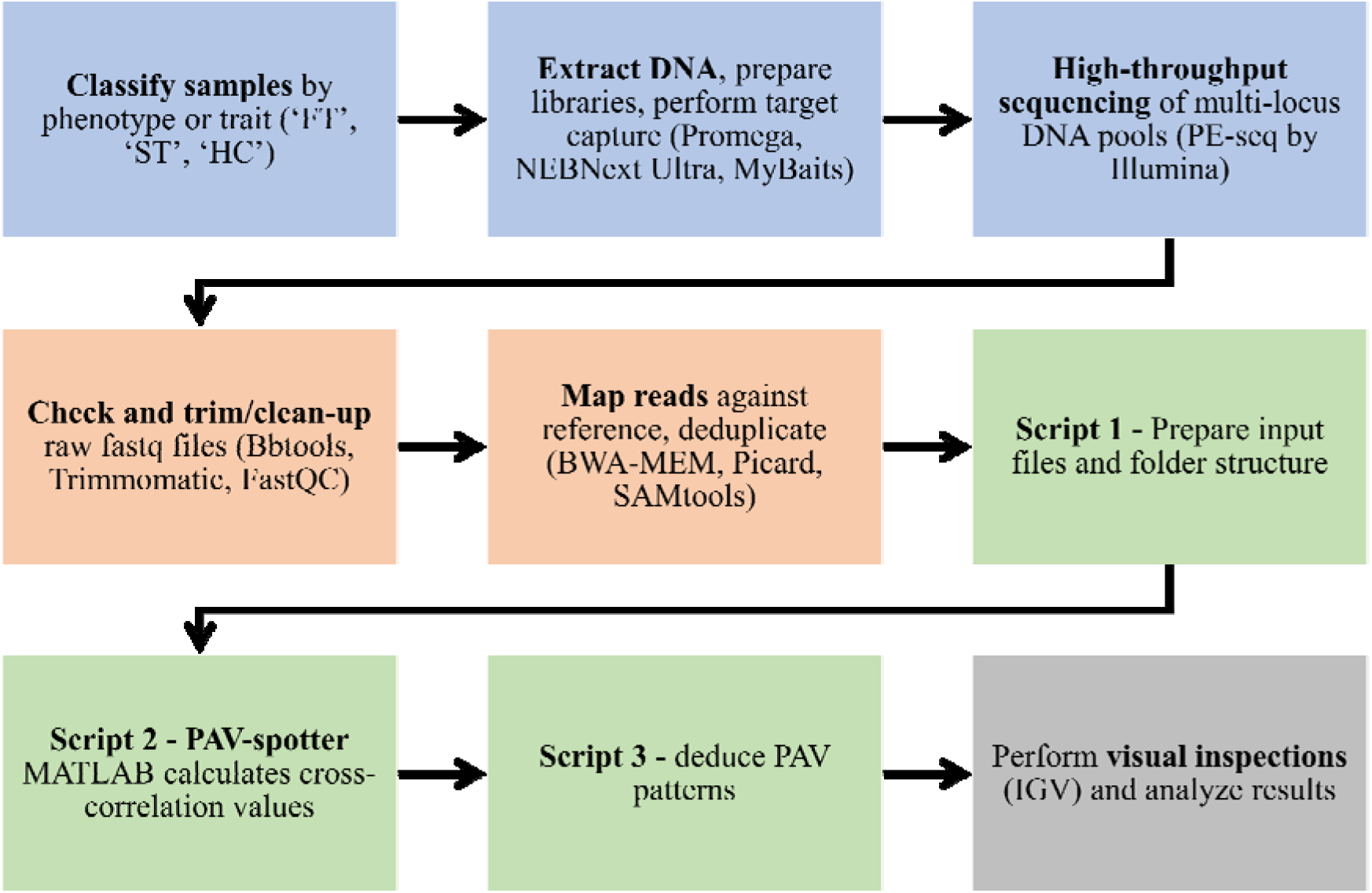
A summary of the consecutive steps of our methods in which the order of the steps is indicated by black arrows. Blue boxes show the laboratory process, orange boxes represent bioinformatic pre-processing steps, green boxes stand for the application of the main PAV-spotter scripts, and the grey box covers the conclusive steps of inspecting and interpreting the results.

Finally, as not much is known about the genetic background of our non-model study species *Triturus*, we performed visual inspections of the read content of all BAM files (n=30) that were used as input for PAV-spotter by checking them in Integrative Genomics Viewer (IGV) software (Robinson et al, 2011) on a Windows environment. This constitutes the last step in our overall workflow (Figure 2). We automated obtaining screenshots through IGV by running batch commands for all 7,139 targets (an example batch script is available as “batch_IGV.bat”, which is written for Windows, but note that the commands used can similarly be run in Unix-based environments such as Linux and macOS). With a ‘checking-by-eye’ approach we categorized PAV hits as ‘likely true’ and ‘likely false’ in order to assess the quantity of false positive outcomes and we cross-checked the results of all the different runs to identify false negative outcomes (in other words: in case a ‘likely true’ target with PAV was retrieved in one analysis, but not in another, we counted it as a false negative in the latter analysis).

We executed all bioinformatic steps, from pre-processing of reads and read depth information to applying PAV-spotter and extracting information from the output, through the High Performance Computing facility ‘ALICE’ (Academic Leiden Interdisciplinary Cluster Environment, the Netherlands). The PAV-spotter GitHub repository provides all scripts and further explanation in the “README.md” file: https://github.com/Wielstra-Lab/PAVspotter.

## Results

A mean of 6,483,062 read pairs were generated on average per sample, with a standard deviation of 1,450,648 read pairs (Supplementary Table 1). After trimming, this changed to a total of 6,187,680 read pairs with a SD of 1,369,254 read pairs. On average, 35.57% of the trimmed reads were successfully mapped against the reference targets after duplicate removal, as an average of 17.09% of all trimmed reads were flagged as duplicates (Supplementary Table 1).

Overall, the targets had a mean read depth of 90.09 sequences and a mean coverage of 97.19 % of the sequence bases (Supplementary Table 2, presented per phenotypic class). For the overall set with ten samples per phenotypic class, the average depth of coverage was 84.7 in the FT group, 97.7 in the group of HC embryos, and 87.8 in the ST embryo group (Supplementary Table 2). Moreover, for the batched samples with five individuals per phenotypic class, this average depth of coverage varied between the lowest number of 76.6 in FT batch 5-2 and the highest number of 102.2 in HC batch 5-1 (Supplementary Table 3). Lastly, for the batched samples with two individuals per phenotypic class, the averages varied between the lowest number of 37.1 in FT batch 2-1, and the highest number of 134.1 in FT batch 2-5 (Supplementary Table 4).

Through the different runs with a sample size of ten per class, we discovered large-scale PAV for in total 72 targets. Genes without PAV had sequence reads with similar distributions for all three classes (Figure 3A), whereas of those 72 aberrant genes that we discovered, 32 showed an absence in FT, but a presence in HC and ST embryos (and are thus “1A-linked”, Table 5 and Figure 3B), while the remaining 40 were absent in ST, but not in HC and FT embryos (and are thus “1B-linked”, Table 6 and Figure 3C). We confirmed our findings using the visual IGV inspection (for examples, see Supplementary Figures 3-5).

**Figure 3:**
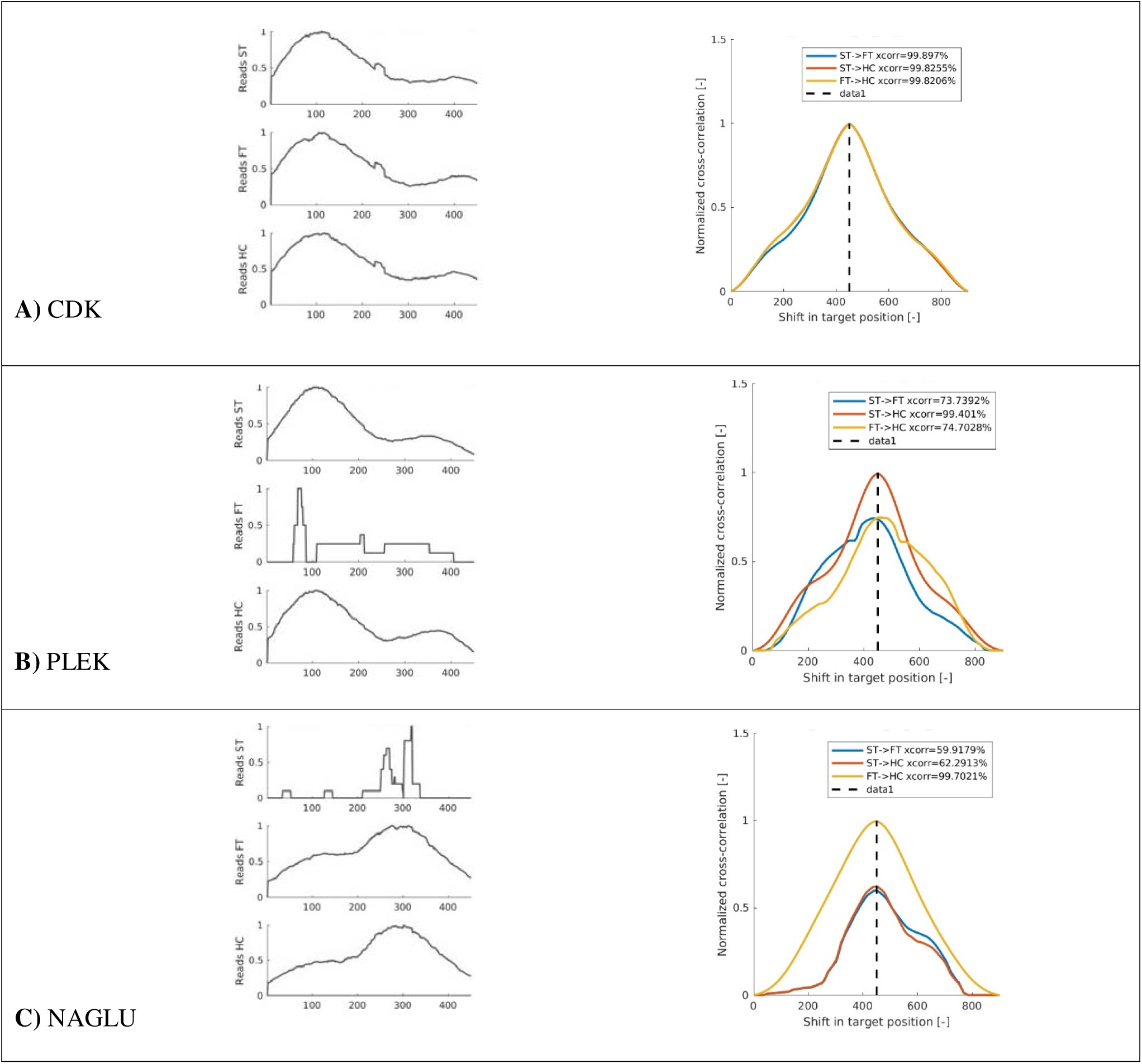
Examples of target-specific plots produced by PAV-spotter in the overall run with no filtering (n=10 per class). The stacked plots on the left side within the panels A-C show the merged and normalized distributions of sequence read depths per phenotypic class, and the colored plots on the right side within the panels A-C show associated measures of similarity of those distributions as a function of the displacement of one relative to the other (including a legenda explaining the colors and cross-correlation values). **A)** An example of a ‘normal’ gene/target, showing the cross-correlation analyses of control marker ‘CDK’ (see Discussion), with a similar shape of the sequence read depth distributions and high correlation values (above the 80% similarity threshold) between all three classes. **B)** An example of a 1A-linked gene/target, showing the cross-correlation analyses of ‘PLEKHM1’ (see Discussion), with a deviant read depth distribution for the FT class and a high correlation value (>80%) for ST vs. HC samples, but a lower cross-correlation value (<80%) for the ST vs. FT and FT vs. HC sample comparisons. **C)** An example of a 1B-linked gene/target, showing the cross-correlation analyses of ‘NAGLU’ (see Discussion), with a deviant read depth distribution for the ST class and a high correlation value (>80%) for FT vs. HC samples, but lower cross-correlation values (<80%) for the FT vs. ST and the ST vs. HC samples.

After running PAV-spotter without any filtering options on the full dataset, we correctly identified all 32 1A targets and generated one false positive in the 1A list, which we identified by the visual IGV inspection (see Supplementary Figure 6). By re-running PAV-spotter with the previously described filtering options, this false positive was removed from the list in some cases, however these additional analyses also generated more false positives and false negatives: the frequency of them depending on the combination of filtering settings used (Supplementary Table 5). For the 40 1B targets, we discovered 37 true positives and three false positives in the unfiltered run. Depending on the combination of filtering settings applied, two of these three false positives again disappeared from the list. However, these extra runs with filtering settings also highlighted three additional 1B targets that were overlooked (as false negatives) in the initial analysis (Supplementary Table 6).

The false positive outcomes consistently had either the lowest - or in a single case, the highest - mean depth values, reflected by the ‘MAXpeak’ output of the overall results matrix generated by PAV-spotter. This value is the peak number of reads in one position of a certain target observed across all the individual samples included in the analyses. The false negative outcomes that came to light as true positive results after additional filtering was applied, all had similarity scores for the resembling classes (FT and HC) above 99% in the unfiltered analysis. But the lower similarity scores of these targets between the non-resembling classes (both between FT and ST and between ST and HC) were lower than 90%, but not lower than the 80% threshold (which is why they were initially overlooked).

The results for the n=5 per class runs (5_1 and 5_2) resemble our earlier findings. We re-discovered an average of 68.5 (95.1% success) out of the total of 72 PAV targets previously discovered, but with half the sample size. For run 5_1, this number was 70 out of 72 PAV targets (97.2% success) and for run 5_2 this number was 67 out of 72 (93.1% success). Overall, for 1A, the results of both the analyses with a sample size of five individuals per class were complementary, as all 32 previously identified 1A targets were re-discovered at least once (Supplementary Table 7). The same goes for the 40 previously identified 1B targets (Supplementary Table 8). Between the two runs, the overall mean depth of coverage was the lowest in 5_2, the analyses that also showed less successful out of the two.

The analysis of the 5_1 subset (n=15) resulted in the discovery of 31 true 1A targets plus one false negative and one two positives, and in 39 true 1B targets with one false negatives and one false positive. The analysis of the 5_2 subset (n=15) again resulted in 31 true 1A targets (with one difference) plus one false negative and two false positives. These false outcomes were not the same as with the 5_1 subset analysis. For 1B, the 5_2 analysis yielded 36 true 1B targets with four false negatives and seven false positives.

The results of the n=2 per class runs (2_1 through 2_5, each with n=6 in total) again highlighted the same PAV exhibiting targets. We re-discovered an average of 70 (97.2% success) out of the total of 72 PAV targets previously discovered, but with a fifth of the sample size (Supplementary Tables 9 and 10). For each of the 2_1, 2_2 and 2_4 runs, these numbers were indeed 70 out of 72 (97.2% success), run 2_3 retrieved 71 out of 72 PAV targets (98.6% success) and run 2_5 69 out of 72 (95.8%).

Overall, for both the 32 previously identified 1A targets and the 40 previously identified 1B targets, these five analyses with a sample size of only two individuals per class appeared complementary, as all true positive targets were re-discovered at least once. For 1A, one of the false negative outcomes came forward as a false negative in two out of the five analyses (and as a true positive in the three other analyses). The other three false negative outcomes in the 1A list were incidental. For 1B, each of the false negative outcomes occurred in only one of the five analyses (and formed a true positive result in the four alternative analyses). The number of false positive outcomes was slightly higher with these low sample size tests (Supplementary Tables 11 and 12), however this was especially noticeable for the 1A results of the first batch (2_1). The mean depth of coverage was also the lowest for the FT (1B1B) samples in this batch (Supplementary Table 4). We therefore tested for a correlation and show that the mean depth of the samples exhibiting absence (i.e. the mean depth of FT samples with determining 1A absence, and the mean depth of the ST with determining 1B absence) appeared to be negatively and significantly correlated to the number of false positives brought forward (Spearmann’s rank correlation, n=10, p=0.018).

## Discussion

We employ a signal cross-correlation approach to discern PAV patterns in target capture data of *Triturus* newt DNA, using transcriptome-based markers with no need for whole genome information. By comparing the read depth in sequence data of embryos of different phenotypic classes, and by manually checking the results of the read alignments, we are able to – within the targeted set of genes – identify over seventy targets that appear to be either present in, or absent from, the genome, depending on the phenotype.

The three example targets displayed in Figure 3 have been independently tested using multiplex (mx) PCR techniques, including mxKASP (Meilink et al, 2025). Control marker CDK is present in all three embryo classes. This corresponds with our results from PAV-spotter, where we discover high cross-correlations values between the distribution of mapped reads of all three phenotypic classes for this target marker (>99% similarity, way above our 80% cutoff threshold). On the other hand, PLEKHM1 is observed in HC and ST embryos, but not in FT embryos, and NAGLU is observed in HC and FT embryos, but not in ST embryos (Meilink et al, 2025). Again, this matches our PAV-spotter findings – even with sample sizes as low as two individuals per phenotypic class.

Evidently, PAV-spotter relies on the correct, a priori classification of samples. When multiple samples are provided, read-depth data are merged per phenotypic class, as the core aim of PAV-spotter is to detect PAV *between* predefined groups rather than *within* them. If desired, the merging and normalization procedure can be disabled to compute cross-correlations among individual samples – for example to explore potential polymorphisms within phenotypic classes. However, this is not the default or recommended use – thus. future users should carefully consider what they want to compare, and why. For instance, we underline that cross-contamination of DNA is a main concern here when using target capture data of ancient DNA (Zavala et al, 2022), as this could potentially distort the similarity values calculated by PAV-spotter – something to be wary of.

The fact that PAV-spotter is able to, on average, re-discover 97,2% of true positive PAV target outcomes in our trials is especially convenient for studies where scientists must rely on a limited amount of available DNA, as is often the case with herbarium specimens (Hart et al, 2016; Kates et al, 2021). However, the highest yield of false positive outcomes is observed in our 1A results of batch 2_1, but not the accessory 1B results, which firstly shows that there is a likely trade-off between sample size and sequence coverage. Preferably, the quality of DNA is as high as possible when working with small sample sizes. Although also preferred in case of a larger sample size per phenotypic class, the chances are then higher that any poor coverage sample(s) will be compensated for by sample(s) with better coverage. In general, spreading sequencing efforts across at least a couple samples, with slightly lower – but still informative – depths per sample, is considered a safer option than working with extremely low sample sizes (Lou et al, 2021; Pezzini et al, 2023).

Regardless of sample size, identifying and characterizing structural variation from target capture data is widely recognized to be difficult (Jones & Good, 2016) and it is especially challenging with non-model organisms. This is because capture-rates may vary considerably depending on bait design, sample quality, species relatedness, batch effects, and stochastic factors (Andermann et al, 2019; Feng et al, 2015). In most cases where targets showed a significant absence of mapped reads, we observe a small amount of reads being (mis)mapped against reference targets when none are expected, for instance (visible in the PAV-spotter output figures and the IGV screenshots). Occasionally, these consist of (clipped/partially matching) reads, something that can be caused by sequencing errors, chimeric reads, errors in the reference sequence, tandem duplications, or genomic rearrangements and structural variants (Schroder et al, 2014; Suzuki et al, 2011). Due to the case-specificity of such potential errors, benchmarking across a broader range of target capture datasets – including those analyzed in independent follow-up studies using PAV-spotter – will be relevant to further assess the accuracy of the PAV-spotter method and, where applicable, to keep building upon its performance over time.

A solution to remove any unwanted (mis-)mapped reads would be to filter more strictly upstream. However, in case of samples or targets that show poor or limited coverage – an issue that is not uncommon with target capture procedures (Bragg et al, 2016) – strict filtering may not be desired. This means that, due to this potential stochasticity, merely using existing tools to check whether there are any mapped reads at all in a sample/target (i.e. checking for the presence of zero reads versus >0 reads), or building *de novo* assembled contigs per target on a sample-per-sample basis, will not be sufficient to identify PAV accurately. Our method offers an alternative solution, as PAV-spotter appears robust enough to detect PAV, even in the face of low coverage and (partially) mis-mapped and clipped reads. However, the trade-off in false positive and false negative outcomes will largely depend on the similarity thresholds and other criteria set by users, as well as on any manual, double-checks performed. General risks associated with target capture experiments, such as inadvertently sequencing paralogues, pseudogenes, or other off-target or multi-copy regions due to probe non-specificity, should therefore always be taken into account: in principle, PAV-spotter is applied under the assumption that targets represent orthologous loci, since potential multi-copy genes were already excluded in the process of designing the target capture probes (De Visser et al, 2025a; Wielstra et al, 2019).

In conclusion, we show that considering the genomic position as a variable for signal displacement instead of time – which is generally the case in more classic cross-correlation applications (Verhaegen & Verdult, 2007) – makes it possible to identify markers of PAV/structural variation in target capture data, without needing any prior knowledge on large-scale, genomic context. And while this tool was specifically tailored for a reciprocal, self-validating test, future users may tweak the code and settings to match their own research intentions. Extensions of the tool may include the incorporation of whole-genome sequence data (Schieffer et al., 2026), or other types of data, to further guide validation and interpretation – meanwhile the current implementation enables robust PAV detection under the constraint of lacking a reference genome, taking away a potential obstacle in study continuations for target capture experts. Our study therefore shows that an innovative and daring, multidisciplinary bioengineering and biotechnological approach can help bioinformatics, and thus the fields of evolutionary and molecular research, forward when dealing with challenging research questions, datasets, and study organisms.

## Acknowledgements, funding, and permits

This project has received funding from the European Research Council (ERC) under the European Union’s Horizon 2020 research and innovation programme (Grant Agreement No. 802759). Sample collection was supported by the Serbian Ministry of Science, Technological Development and Innovation (grants nos. 451-03-66/2024-01/200007, 451-03-65/2024-03/200178, 451-03-66/2024-03/200178). Dr. Ana Ivanović supported breeding and collecting *Triturus* samples. Individuals of *T. ivanbureschi* and *T. macedonicus* were originally collected from natural populations: *T. ivanbureschi* from Zli Dol, Serbia with permission obtained from the Serbian Ministry of Energy, Development and Environmental Protection (permit No. 353-01-75/2014-08) and *T. macedonicus* in Ceklin, Montenegro with permission obtained from the Agency for Environmental Protection, Montenegro (permit No. UPI-328/4). The experimental procedures were approved by the Ethics Committee of the Institute for Biological Research „Siniša Stanković”, University of Belgrade (decisions no. 03-03/16 and 01-1949). All experiments were performed in accordance with the Directive 2010/63/EU.

## Data Availability and Benefit-Sharing

The raw, Illumina sequencing reads used in this study have been submitted to the NCBI Sequence Read Archive (SRA) and are publicly available through BioProject PRJNA1111729 (https://www.ncbi.nlm.nih.gov/sra/PRJNA1111729). The PAV-spotter tool, including further explanation on how to use and customize the underlying code, is available through the PAV-spotter GitHub repository: https://github.com/Wielstra-Lab/PAVspotter. Furthermore, Supplementary Materials are provided through a Zenodo repository: https://zenodo.org/records/13991751. Figures created by PAV-spotter for each separate analysis that are not shown in the paper can be provided upon request. The same goes for all IGV screenshot images generated by the batch script.

## Author contributions

MdV and CvdP designed the tool. BW, MdV and CvdP designed the experiments. MC, TV and AI collected the samples and conducted the phenotypic classification. MdV and AT performed the molecular laboratory work. CvdP wrote the PAV-spotter scripts (Scripts 2a and 2b, both the MATLAB and the Python versions), which comprises of the signal cross-correlation code. Other scripts (Script 1, Script 3, and the IGV batch script) were written by MdV. MdV conducted the main bioinformatics, data acquisition, data analyses, and data interpretation. MdV drafted the work and CvdP added mathematical details to the text and conducted p-value/threshold estimations. All authors revised the manuscript and approved of the submitted version.

## Declaration of Generative AI Use

Generative AI tools, such as ChatGPT, were used for minor linguistic refinements, including occasional adjustments to word choice or sentence structure in written text. Generative AI was, however, not used to conduct the research, perform the data analyses, or generate the original scientific study design or content – including the resulting manuscript text, images, or figures. The only exception regarding code development was the use of ChatGPT to assist in translating originally human-written MATLAB code into Python language (followed by manual testing and optimization).

## Competing Interests

The authors declare that they have no competing interests.

---

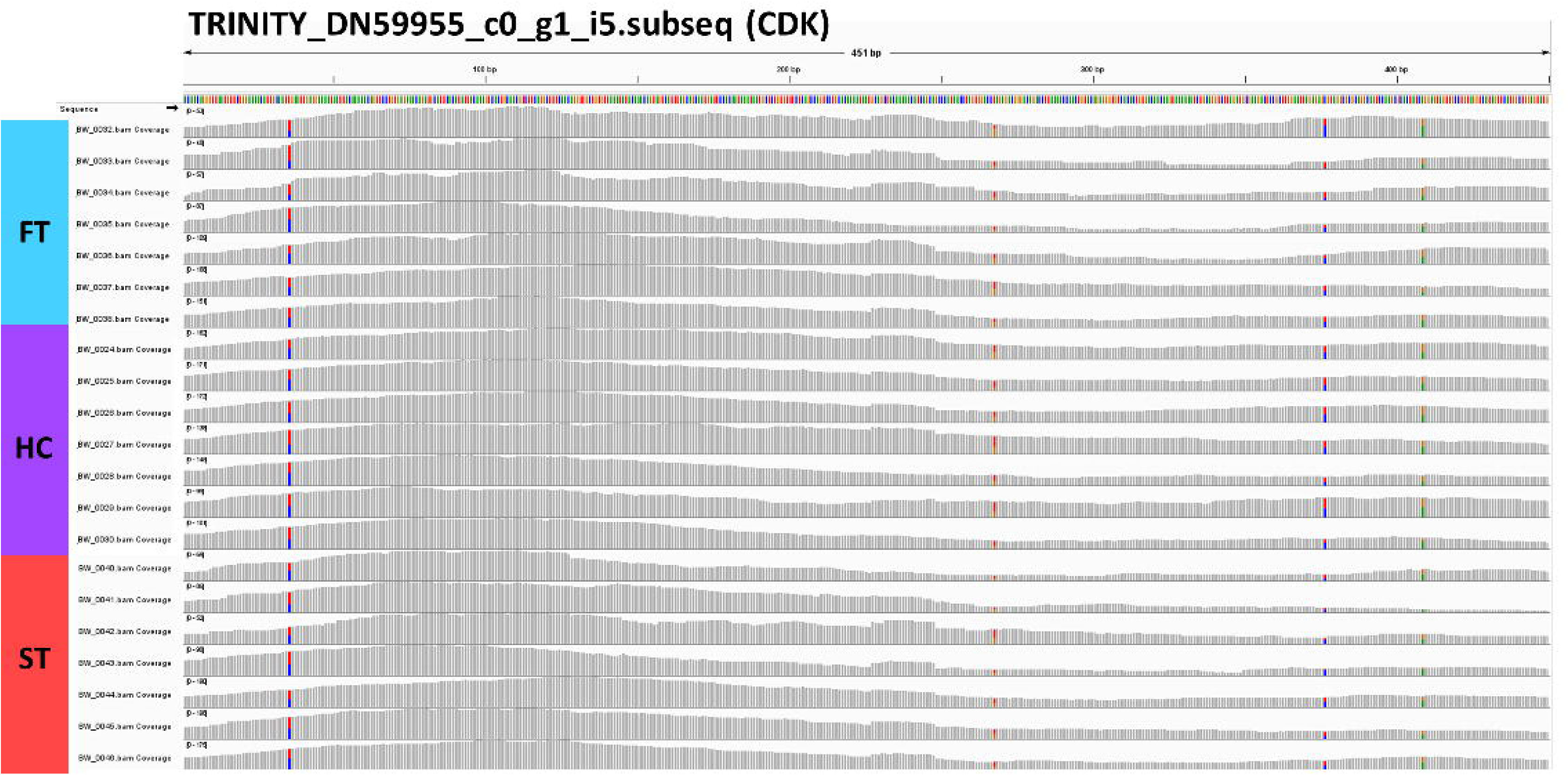
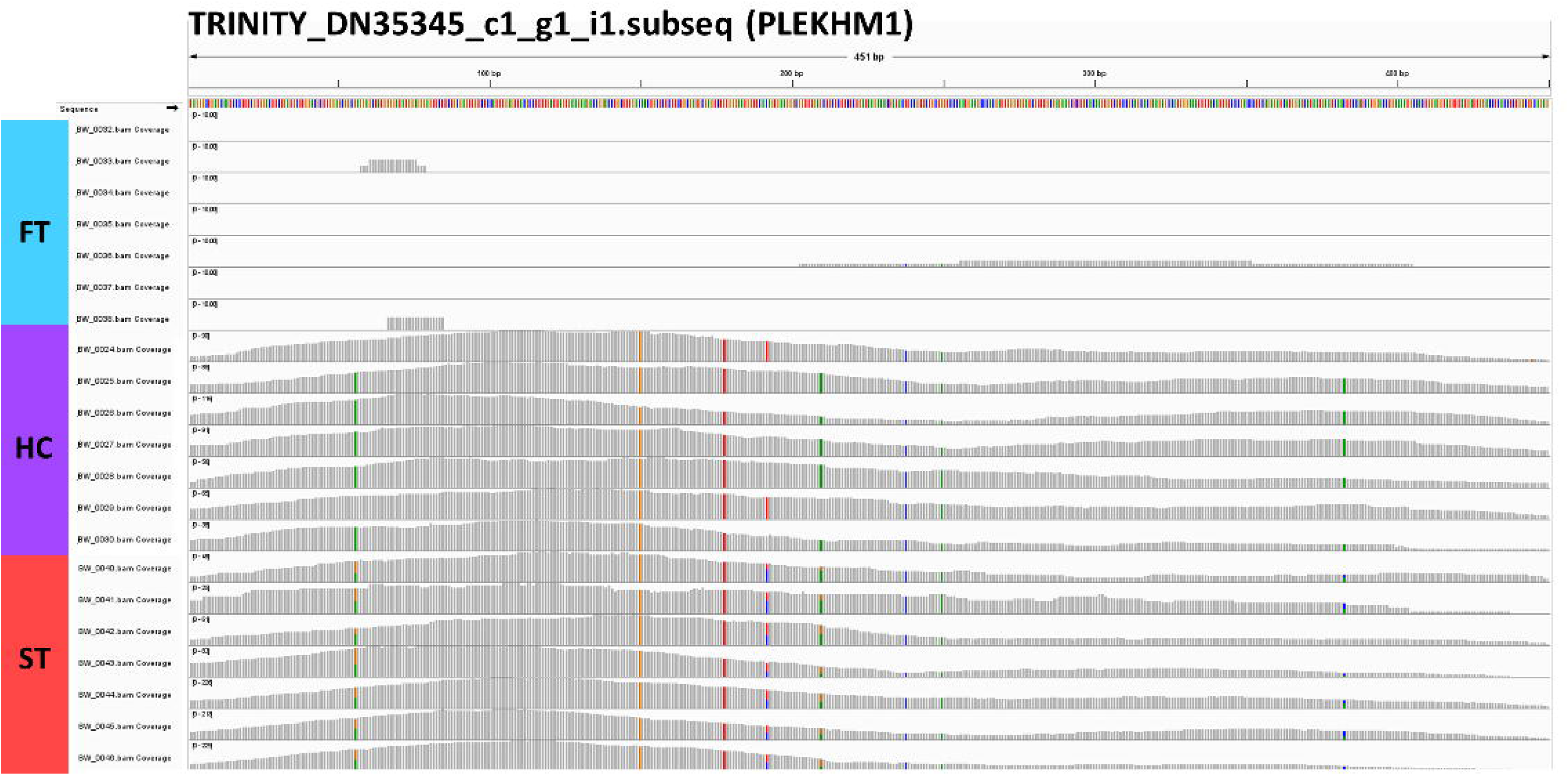
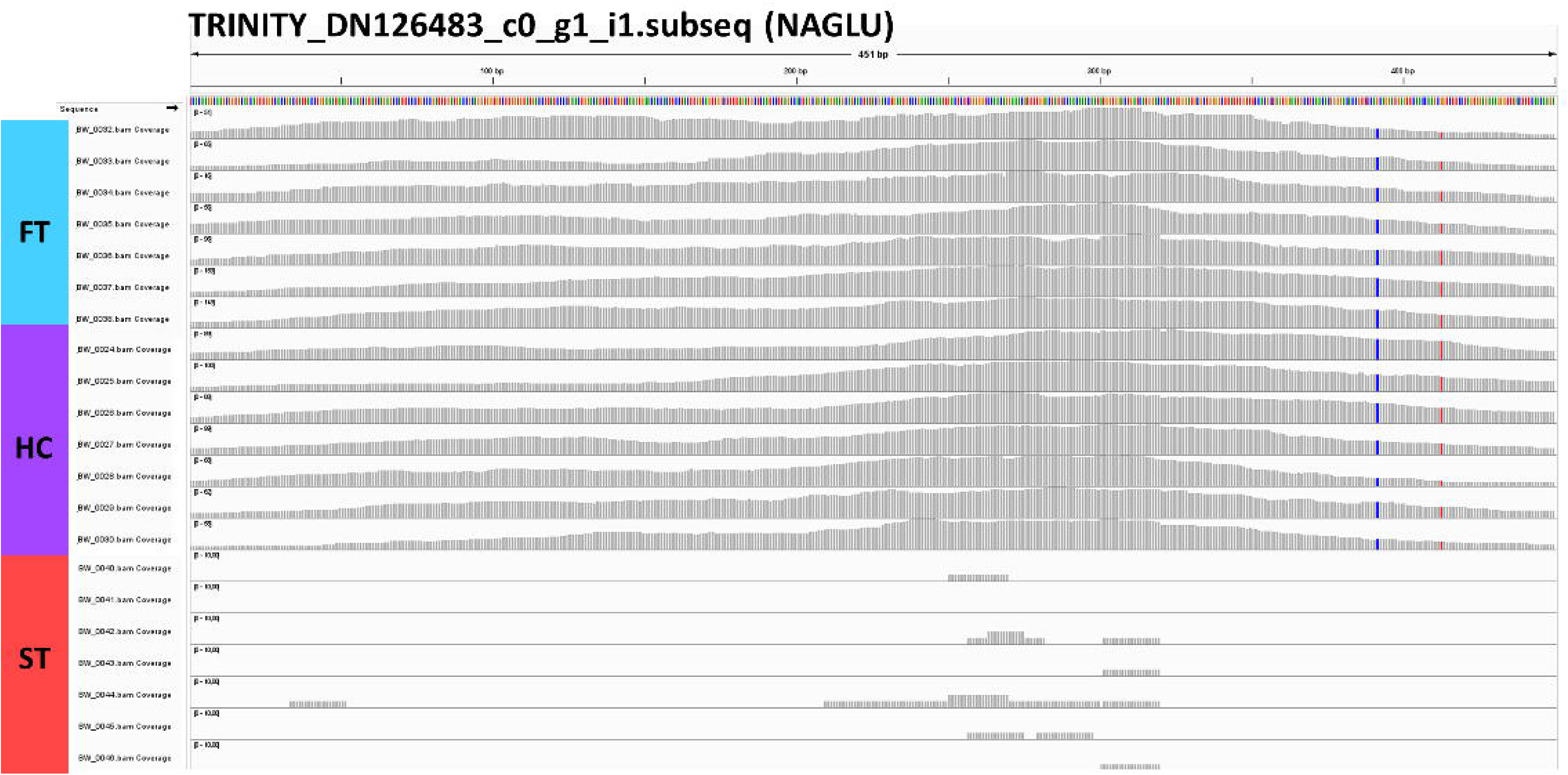
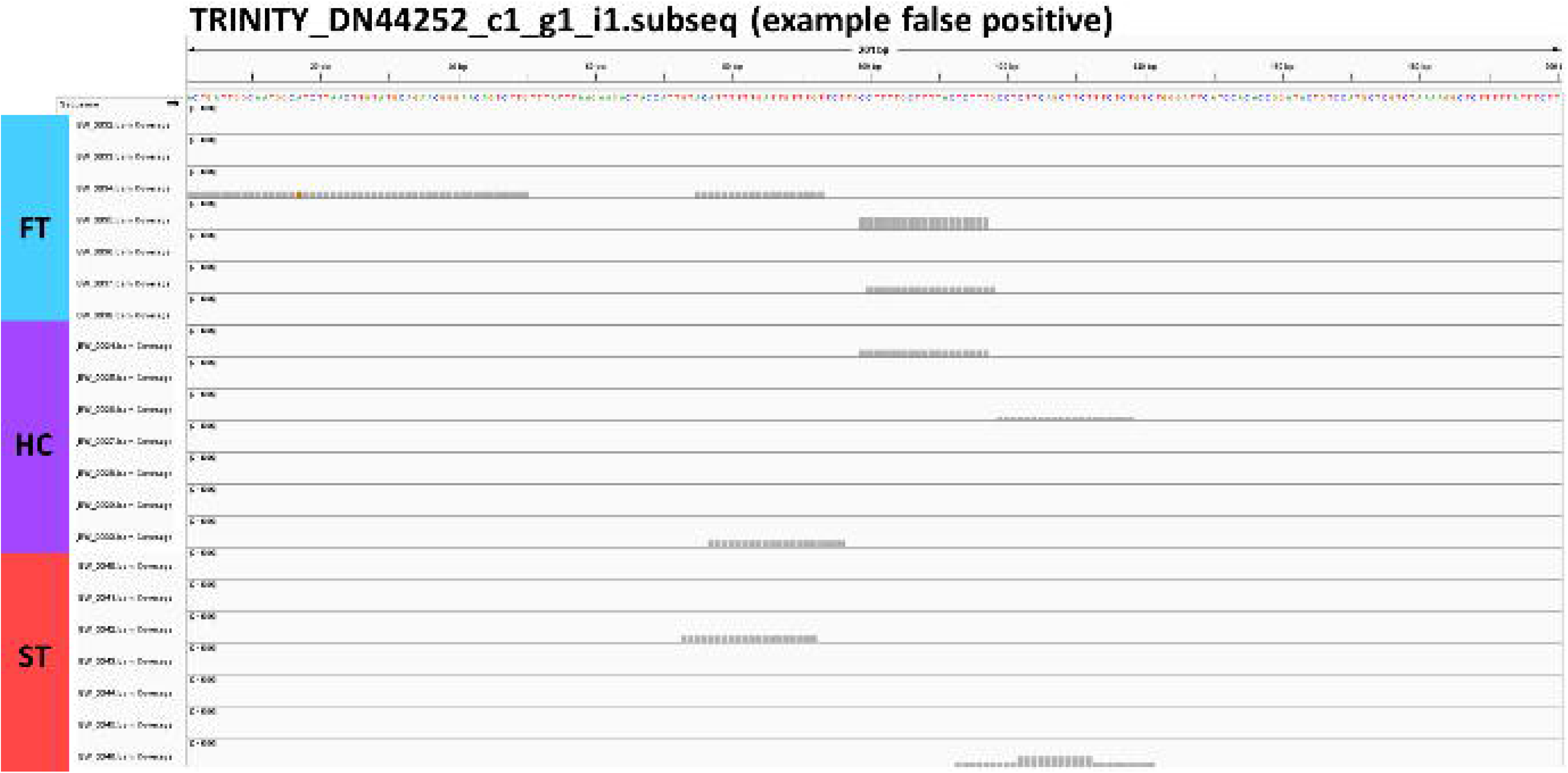
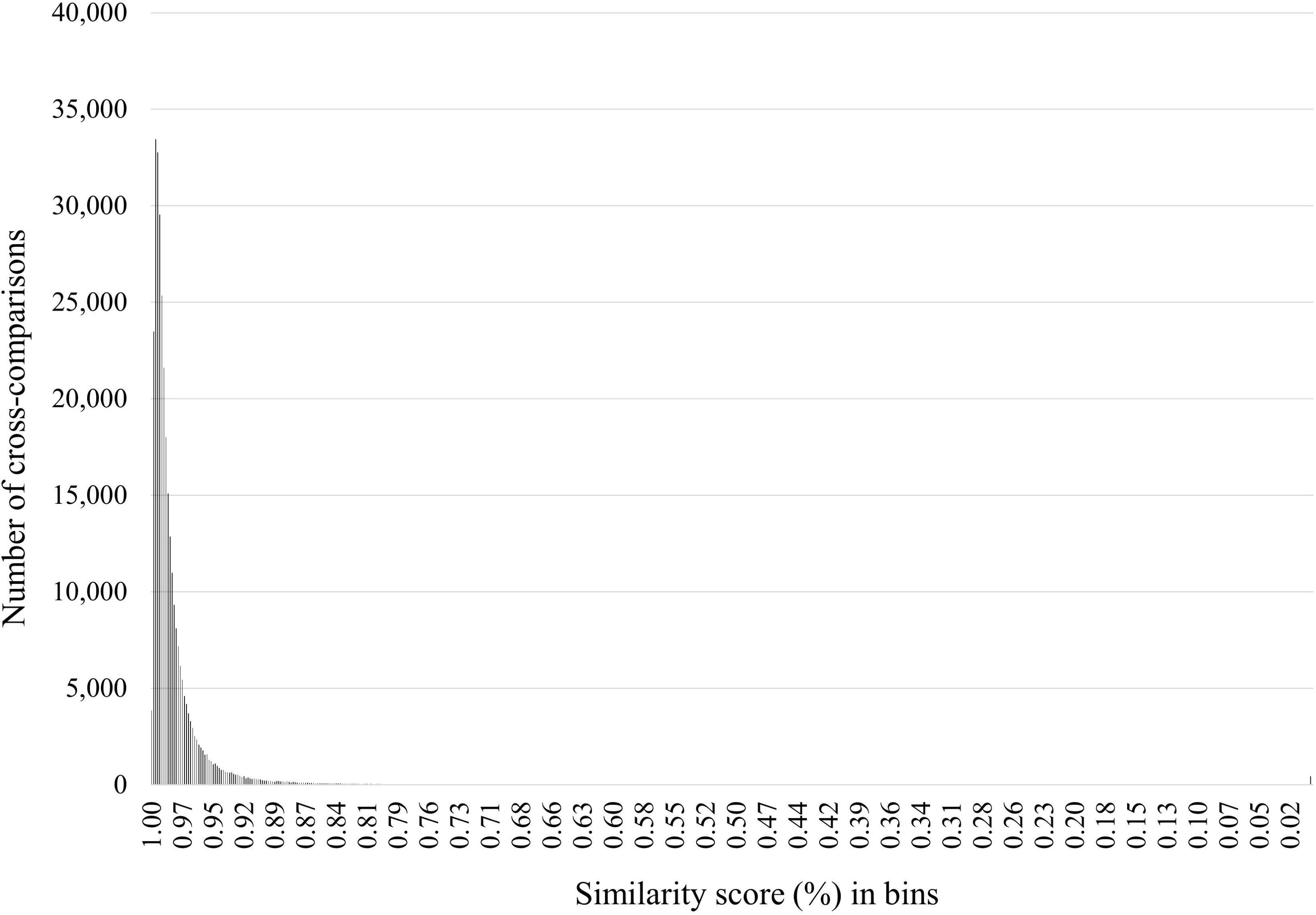
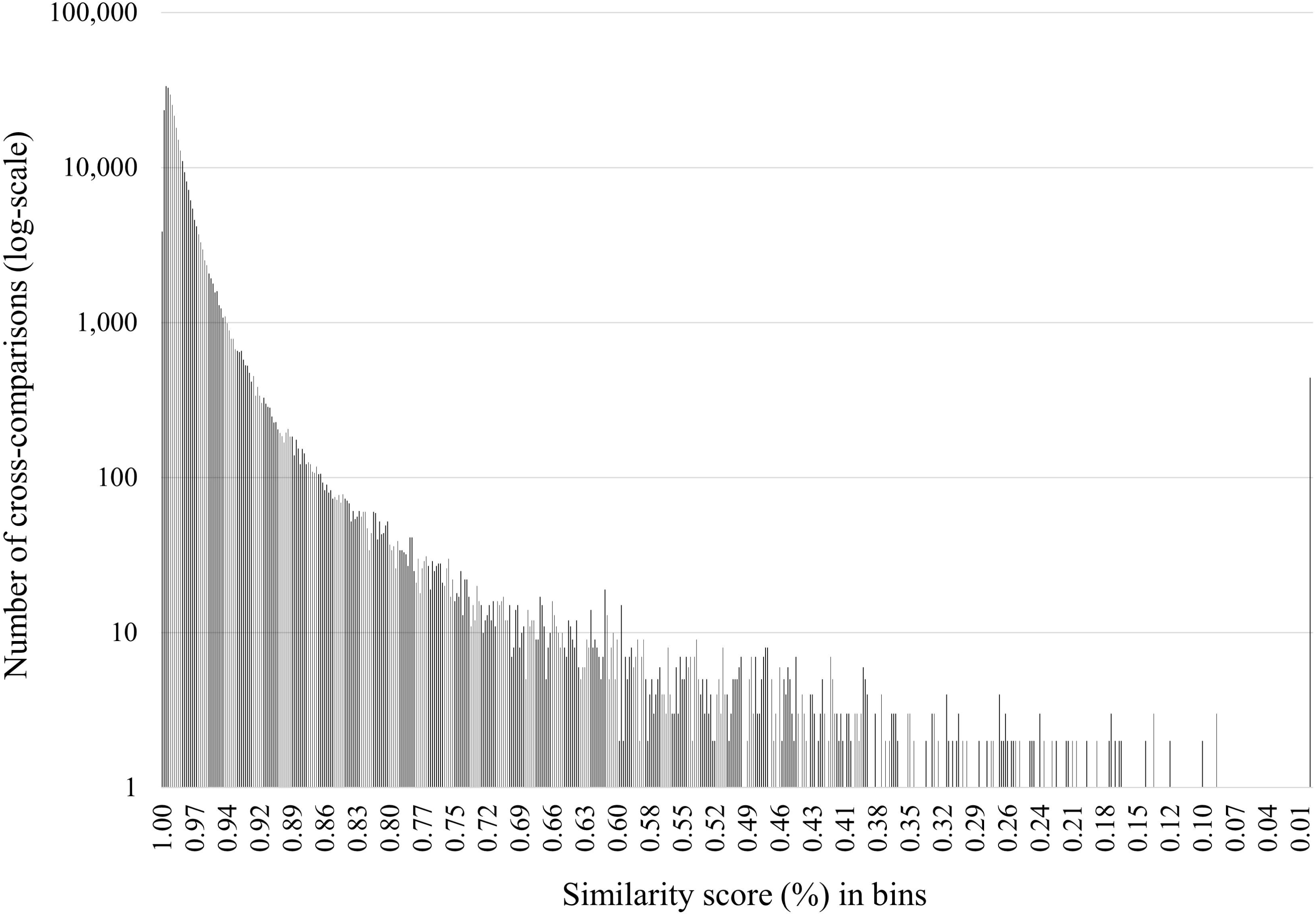

## Notes

https://github.com/Wielstra-Lab/PAVspotter

https://www.ncbi.nlm.nih.gov/sra/PRJNA1111729

https://zenodo.org/records/13991751

### Competing Interest Statement

The authors have declared no competing interest.

### Summary of Updates

We have updated some of the text and headers, but the concept, experiments, analyses and results are remain the same. They are minor tweaks and nothing about the study or contents itself has changed inherently. Also: two additional sources/references have been added to the article (both from 2026).

